# N-type calcium channels control GABAergic transmission in brain areas related to fear and anxiety

**DOI:** 10.1101/2020.05.24.113712

**Authors:** Maxwell Blazon, Brianna LaCarubba, Alexandra Bunda, Natalie Czepiel, Shayna Mallat, Laura Londrigan, Arturo Andrade

## Abstract

Presynaptic N-type (Ca_v_2.2) calcium channels are key for action potential evoked transmitter release in the peripheral and central nervous system. Previous studies have highlighted the functional relevance of N-type calcium channels at both the peripheral and central level. In the periphery, the N-type calcium channels regulate nociceptive and sympathetic responses. At the central level, N-type calcium channels have been linked to aggression, hyperlocomotion, and anxiety. Among the areas of the brain that are involved in anxiety are the basolateral amygdala, medial prefrontal cortex, and ventral hippocampus. These three areas share similar characteristics in their neuronal circuitry. Pyramidal projection neurons are under the inhibitory control of a wide array of interneurons including those that express the peptide cholecystokinin. This type of interneuron is well-known to rely on N-type calcium channels to release GABA in the hippocampus, however whether these channels control GABA release from cholecystokinin-expressing interneurons in the basolateral amygdala and medial prefrontal cortex is not known. Here using sophisticated mouse models to genetically label cholecystokinin-expressing interneurons, we found that N-type calcium channels control ~50% of GABA release in basolateral amygdala. By contrast, N-type calcium channels are functionally absent in synapses of cholecystokinin-expressing interneurons of medial prefrontal cortex. Our findings provide insights into the precise localization of N-type calcium channels in synapses of brain areas related to anxiety.

## 1. Introduction

Presynaptic N-type (Ca_v_2.2) calcium channels open in response to depolarization induced by action potentials, thereby contributing to neurotransmitter release. N-type calcium channels, together with P/Q-type (Ca_v_2.1) and R-type (Ca_v_2.3) calcium channels, mediate neurotransmission in peripheral and central synapses [1–3]. In the periphery, N-type calcium channels are essential for the release of neurotransmitters from nociceptors onto neurons of the dorsal horn of the spinal cord, thus controlling the transmission of noxious information to the central nervous system (CNS) [4]. Furthermore, N-type calcium channels control the release of adrenaline from postganglionic sympathetic neurons [5–8]. In the central nervous system, N-type calcium channels are localized to specific synapses, thus their contribution to transmitter release is cell-specific. Dopaminergic neurons of the substantia nigra and ventral tegmental area heavily rely on N-type calcium channels to release dopamine [9–11]. In the hippocampus (HPC), interneurons that express the peptide cholecystokinin (CCK^+^INs) fully rely on N-type calcium channels to release GABA [12–16]. This is in contrast to interneurons that express somatostatin (SOM) and parvalbumin (PV), which utilize P/Q-type calcium channels to release GABA [13, 17, 18]. In terminals of pyramidal projection neurons (PNs), N-type calcium channels work together with P/Q- and R-type calcium channels to regulate glutamate release [19–21]. Due to this broad expression, disruption in N-type calcium channel activity leads to alterations of peripheral and central nervous system functions.

Behavioral studies in the Ca_v_2.2-null mice highlight the functional role of N-type calcium channels in regulating processes related to the peripheral and central nervous system. Regarding the peripheral nervous system, defects in sensory and sympathetic functions have been observed in Ca_v_2.2-null mice. In peripheral sensory processing, Ca_v_2.2-null mice show reduced thermal nociceptive responses compared to WT mice [22]. Ca_v_2.2-null mice also exhibit reduced thermal hyperalgesia in inflammatory pain models [23, 24]. In sympathetic function, Ca_v_2.2-null mice show elevated heart rate and blood pressure compared to WT mice [7]. At the central level, Ca_v_2.2-null mice exhibit hyperlocomotion, enhanced aggression, enhanced response to apomorphine, reduced exploratory behavior, and increased freezing during startle [23, 25, 26]. The latter two observations implicate N-type calcium channels in anxiety-related behaviors.

Additional evidence implicating N-type calcium channels in anxiety arises from behavioral analysis of novel mouse genetic models, behavioral studies with brain infusions of ω-conotoxin GVIA (ω-ctx GVIA), and clinical studies in humans. Mice with restricted splice choice in the gene that encodes for the α_1_-pore forming subunit of Ca_v_2.2, *Cacna1b* (lack *37a-Cacna1b* splice variant), show alterations in exploratory behavior in basal conditions and under mild stress [27]. Infusions of ω-ctx GVIA in the medial prefrontal cortex (mPFC) impairs recall of extinguished memories in fear conditioning, and ventricular application of ω-ctx GVIA results in reduced exploratory behavior in the elevated plus maze and light-dark box [28, 29]. Increase in anxiety has been observed in individuals undergoing treatment for chronic pain using the N-type calcium channel blocker Ziconotide [30]. All of these observations suggest that the N-type calcium channels can potentially serve as a target to treat anxiety disorders. In conjunction with this, the N-type calcium channel blockers CNV2197944 and Z160 are currently being tested or have been tested in clinical trials for anxiety [31]. However, few studies exist determining the functional expression of N-type calcium channels in areas related to anxiety disorders such as the basolateral amygdala (BLA) and mPFC.

The BLA and mPFC are well-known brain areas to encode fear and anxiety in humans and rodents [32]. In rodent models, the BLA and mPFC regulate anxiety-and fear-related behavior in opposing ways [33–36]. Direct activation of the BLA leads to reduced exploratory behavior in the elevated plus maze and enhanced freezing in response to auditory cues initially paired to a shock [37]. By contrast, activation of the mPFC results in enhanced exploratory behavior and reduced freezing in fear conditioning tasks [36].

The BLA and mPFC contain similar cell-types. In both areas, several types of interneurons, distinguished by the expression of peptides, have a powerful inhibitory control on PNs. Among the main interneuron types that control PN activity are interneurons that express PV (PV^+^INs), SOM (SOM^+^INs), vasoactive intestine peptide (VIP^+^INs), and CCK^+^INs [38, 39]. PV^+^INs synapse onto cell bodies; CCK^+^INs synapse onto cell bodies and dendrites; and SOM^+^INs synapse onto dendrites of PNs [40–42]. VIP^+^INs primarily suppress SOM^+^INs, thereby disinhibiting PNs [43].

Previous studies have shown that CCK^+^INs play important roles in fear-and anxiety-related behavior [44]. Selective activation of CCK^+^INs enhances contextual fear conditioning [45]. CCK^+^INs mediate the effects of endocannabinoids on anxiety and stress [46]. Optogenetic activation of CCK^+^INs of the BLA *in vivo* facilitates fear extinction [47]. Here, we determined whether N-type calcium channels control GABA release from CCK^+^INs in the BLA and mPFC. We used genetic mouse models, electrophysiology, and pharmacology to specifically label CCK^+^INs, as well as to record GABA release from CCK^+^IN synapses onto PNs in the BLA and mPFC. We found that CCK^+^INs in the BLA partially rely on N-type calcium channels to release GABA, whereas CCK^+^INs in mPFC lack functional N-type calcium channels. However, N-type calcium channels partially control GABA release from other interneuron types in the mPFC. These observations suggest that N-type calcium channels have a cell- and region-specific role on GABA release. This is significant because determining the functional expression of N-type calcium channels in circuits related to fear and anxiety could help strengthen the links between N-type calcium channels and anxiety.

### Ethics Statement

All procedures were approved by the Institutional Animal Care and Use Committee at the University of New Hampshire.

## 2. Materials and Methods

### 2.1. Animal Models

Adult female or male mice were used in all of our experiments. No differences associated with sex were detected in our experiments. To visualize CCK^+^INs in the BLA and mPFC, we utilized intersectional labeling with Cre and Flpe recombinases under the CCK and Dlx5/6 promoters, respectively [48, 49]. We used this approach because CCK is broadly expressed in both PNs and interneurons, whereas Dlx5/6 is restricted to interneurons of the forebrain (**Figure 1**) [48, 50]. We first crossed *Cck-IRES-Cre* mice (The Jackson Laboratory, 012706) with *Dlx5/6-Flpe* mice (The Jackson Laboratory, 010815). The progeny from this initial cross, *Cck-IRES-Cre; Dlx5/6-Flpe (CCK-Dlx5/6),* were dual transgenic mice with both alleles. Next, we crossed these mice with *Ai65(RCFL-tdT-D)* mice (The Jackson Laboratory, 021875). *Ai65(RCFL-tdT-D)* mice express the red fluorescent protein tdTomato (tdT) under the control of two STOP codons. The first STOP cassette was flanked by loxP sites (recognized by Cre) and the second was flanked by FRT sites (recognized by Flpe). The progeny from this cross was triple-transgenic mice with the genotype *Cck-IRES-Cre; Dlx5/6-Flpe; Ai65(RCFL-tdT)-D (CCK-Dlx5/6-tdT)* (**Figure 1A**). In these mice, Cre-Lox and Flpe-FRT recombination removed the two STOP cassettes, resulting in tdT expression in CCK^+^INs. Successful labeling of CCK^+^INs was validated in a previous work from our lab [27].

**Figure 1.**
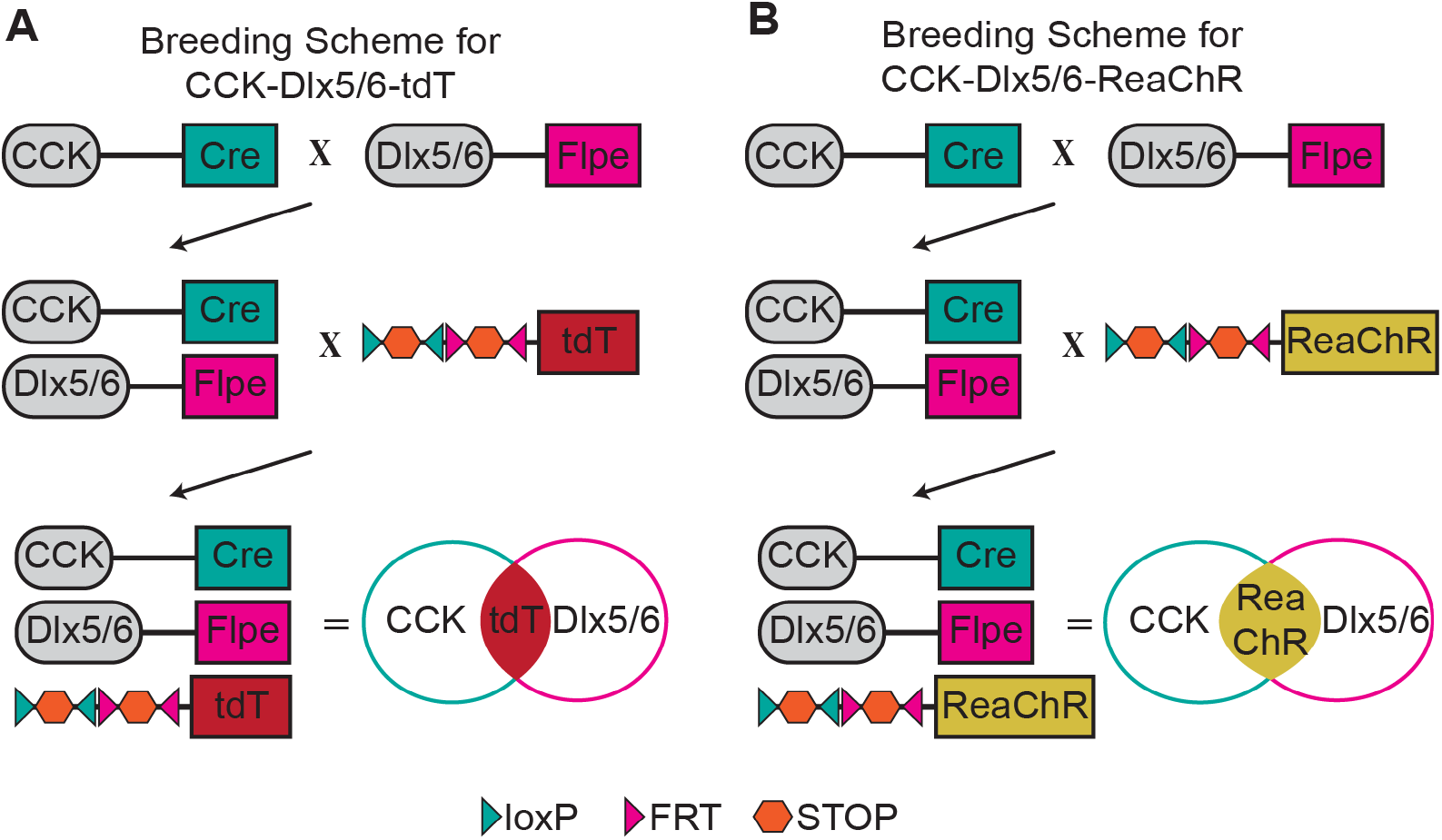
Breeding scheme to generate mice expressing tdT and ReaChR in CCK^+^INs for confocal microscopy and electrophysiology, respectively. **A**) CCK-Cre mice were bred with Dlx5/6-Flpe mice to generate dual transgenic mice (CCK-Dlx5/6), which were crossed with Ai65 mice containing a tdT allele preceded by two STOP codons. Two loxP sites flank the first STOP codon and FRT sites flank the second STOP codon. The offspring of these mice containing the three alleles (CCK-Dlx5/6-tdT) resulted in expression of tdT in CCK^+^INs. **B**) To record synapses of CCK^+^INs onto PNs, we generated triple transgenic mice that expresses ReaChR in CCK^+^INs and applied optogenetics. Here the dual transgenic mice CCK-Dlx5/6 were bred with R26 LSL FSF mice, which contain the allele ReaChR preceded by two STOP codons similar to Ai65 mice. Therefore, after removal of the two STOP codons by Cre and FLPe, ReaChR was expressed in CCK^+^INs.

To perform electrophysiological studies of synapses between CCK^+^INs and PNs, we used optogenetics. CCK^+^INs were labeled with red-shifted channelrhodopsin (ReaChR) fused to m-Citrine. To accomplish this, we generated a second triple transgenic mouse line (**Figure 1B**). We crossed *CCK-Dlx5/6* mice to R26 mice *(R26 LSL FSF ReaChR-mCitrine,* The Jackson Laboratory, 024846), which expressed ReaChR under the control of two STOP codons flanked with loxP and FRT sites. The progeny from this cross are triple-transgenic mice with the genotype *Cck-IRES-Cre; Dlx5/6-Flpe; R26 LSL FSFReaChR-mCitrine (CCK-Dlx5/6-ReaChR).*

### 2.2. Genotyping

Mice were genotyped using toe biopsy as done previously [27]. Primer sequences and band size are located in **Table 1.**

**Table 1.**
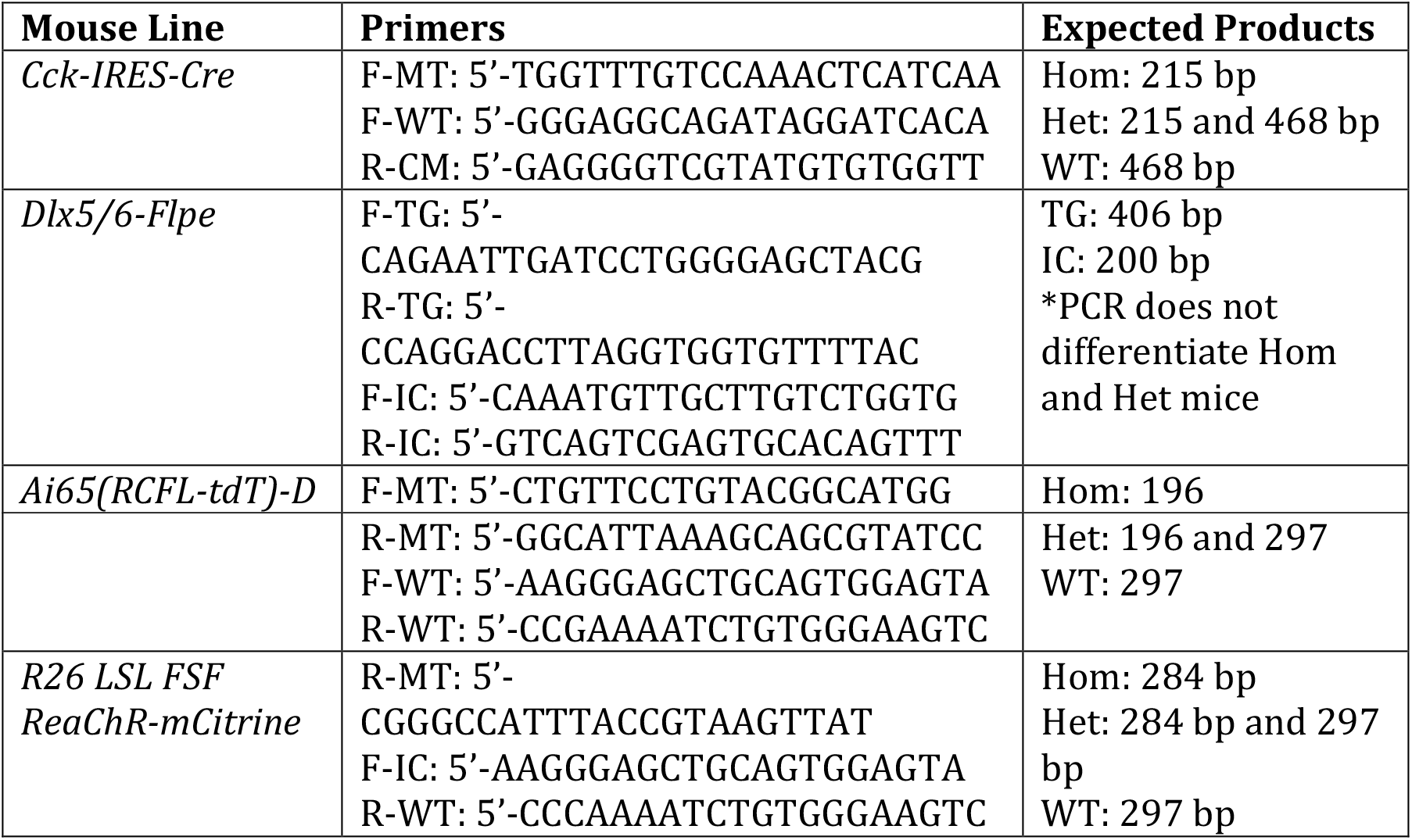
Primer sequences and band sizes for mouse genotyping. Information on the primer sequences and expected PCR products for each mouse line used. F, forward primer sequence; R, reverse primer sequence; MT, mutant; WT, wild-type; CM, common primer; Hom, homozygous; Het, heterozygous; TG, transgene; IC, internal control.

### 2.3 Histochemistry and confocal microscopy

Adult *CCK-Dlx5/6-tdT* mice were deeply anesthetized with intraperitoneal injections of EUTHASOL^®^ (710101, Virbac Co.). Cardiac perfusions were performed using formalin:PBS solution (HT501128, Sigma-Aldrich). Brains were rapidly dissected and stored at 4°C in formalin solution for 24-72 hours for further fixation. 100 μm coronal brain slices were prepared using a vibratome (VT1000 S, Leica) and transferred to a 12-well plate (Greiner Bio-One). Slices were washed for ten minutes three times in PBT (0.2% Triton X, T8787; Sigma-Aldrich and PBS, P3813; Sigma-Aldrich) on an orbital rocker (SK-O180-S, Scilogex). To identify cell bodies, we stained nucleic acids with SYTO-13 (S7575, ThermoFisher) by incubating slices for one hour at room temperature in a 1:10000 dilution of SYTO-13 in PBT. Following incubation, slices were washed twice in PBT and once in PBS before mounting. For brain slices with multiple fluorescent proteins, multi-track acquisition was implemented to avoid excitation cross talk. Fluorophores and their spectra were as follows (excitation/emission in nm): SYTO-13 (488/509), mCitrine (514/529), and tdTomato (554/581). Images were acquired using a Zeiss LSM 510 Meta and the proprietary software.

### 2.4. *In Vitro* Brain Slice Preparation

Adult *CCK-Dlx5/6-ReaChR* mice were deeply anaesthetized with isoflurane and quickly decapitated. Brains were rapidly removed from the skull and placed into chilled and oxygenated (95%_O_2/CO_2_) artificial cerebral spinal fluid (cutting aCSF) containing (in mM): NaCl (130), KCl (3.5), KH_2_PO_4_ (1.1), MgCl_2_ (6), CaCl_2_ (1), dextrose (10), kynurenic acid (2), NaHCO3 (30), ascorbate (0.4), thiourea (0.8), and sodium pyruvate (2) at pH 7.35 and 310 mOsm. 300 μm coronal slices containing the BLA or mPFC were prepared using the vibratome Leica VT1200 S (Leica Biosystems). Slices were transferred from the vibratome to a brain slice chamber (BSKH; Digitimer) containing oxygenated cutting aCSF and kept at 37°C for 15 minutes before transferring to a second slice chamber containing oxygenated room temperature recording aCSF that contained (in mM) NaCl (130), KCl (3.5), KH2PO4 (1.1), MgCl_2_ (1.3), CaCl_2_ (2.5), dextrose (10), NaHCO3 (30), ascorbate (0.4), thiourea (0.8), and sodium pyruvate (2) at pH 7.35 and 300 mOsm. Slices were allowed to stabilize for one hour before transferring to the recording chamber where they remained for no more than two hours during patch-clamp recordings.

### 2.5. Electrophysiology

Whole-cell voltage and current clamp recordings were performed on visually identified PNs of the BLA and mPFC. Cells were identified in the brain slice with differential interference contrast (DIC) microscopy using a camera (01-ROL-BOLT-M-12, QImaging) mounted on an upright microscope (BX51WI; Olympus). All recordings were performed with a 700B amplifier (Molecular Devices) and digitized with a 1550A analogue/digital convertor (Molecular Devices). Data were acquired at 20 KHz and filtered at 2 KHz with pCLAMP 10 (Molecular Devices). Brain slices were perfused with recording aCSF at 1-2 mL/min with constant oxygenation. Borosilicate glass micropipettes with tip resistances between 3-5 MOhms were used to perform whole-cell current or whole-cell voltage clamp experiments. To record action potentials we used the following intracellular solution (in mM): K-gluconate (140), HEPES (10), MgCl_2_ (3), K-ATP (2), Na_2_GTP (0.4), and phosphocreatine (5) at pH 7.4 and 290 mOsm. To record inhibitory postsynaptic currents (IPSCs) we utilized a cesium-based intracellular solution consisting of (in mM): Cs-gluconate (140), HEPES (10), MgCl_2_ (3), K-ATP (2), Na_2_GTP (0.4), and phosphocreatine (5) at pH 7.4 and 290 mOsm. PNs in the BLA and mPFC were identified by their pyramidal-like morphology. To evoke action potentials, several square pulses of positive current were injected into the cells. For IPSC recordings, PNs were patched and held at −70 mV. IPSCs were evoked with LED light filtered with Texas Red filter. LED stimuli was applied for 0.1 ms every 10 s. Pulses of red light were controlled with a digital input of a 1550A analogue/digital convertor. Series resistance was measured throughout the experiment and cells with an increase in >10% of series resistance were discarded. For electrically evoked IPSCs, an electrical current was delivered to brain slices using a current stimulus isolator (A365, WPI) coupled to a tungsten concentric electrode (FHC). Pulses of 0.1 ms were delivered every 10 sec using a digital input of a 1550A analogue/digital convertor. Current intensity was adjusted to obtain an IPSC amplitude between 300 and 500 pA, thereby reducing voltage errors.

### 2.6. Pharmacology

Stock concentrations were made at 1000x working concentrations and stored at – 20°C. 100 μM BIC (ab120107; Abcam) was added to the bath perfusion to confirm IPSCs. 0.5 μM ω-ctx GVIA (ab120215, Abcam) was added to the bath perfusion. 50 μM APV (ab120030; Abcam) and 20 μM NBQX (ab120046, Abcam) was added to the bath to block glutamatergic neurotransmission when electrical stimulation was used. Aliquots were prepared for each stock solution and thawed only once on the day of recordings.

## 3. Results

We first confirmed that CCK^+^INs were present in the BLA and mPFC in our CCK-Dlx5/6-tdT mice. We performed immunohistochemistry in sections of both the BLA and mPFC of CCK-Dlx5/6-tdT mice. The identity of these cells was previously confirmed using fluorescence activated cell sorting coupled to qRT-PCR. Cells expressing tdT also expressed GABAergic markers such as glutamate decarboxylase-2 mRNA and the cannabinoid receptor 1 mRNA; both are hallmarks of CCK^+^INs [27]. As expected, we found cells labeled with tdT in both areas. CCK^+^INs were scattered throughout the BLA (**Figure 2A**). Similarly, CCK^+^INs were found distributed across the layers I-VI of the mPFC (**Figure 2B**). These results suggest that we successfully labeled CCK^+^INs in BLA and mPFC using an intersectional genetic labeling.

**Figure 2.**
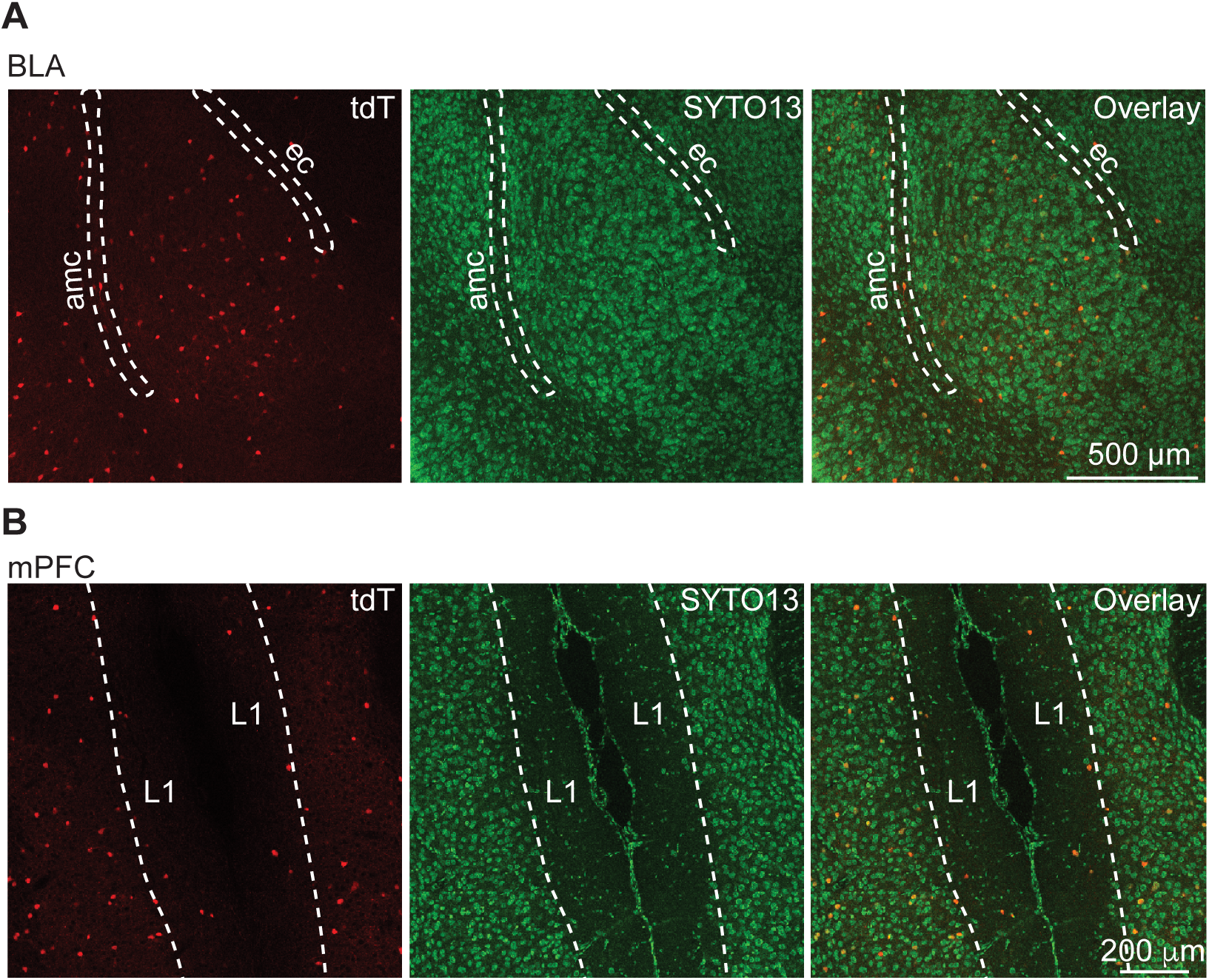
Presence of CCK^+^INs in the BLA and mPFC. **A**) Representative fluorescence images from the BLA of a CCK-Dlx5/6-tdT mouse. CCK^+^INs (red) were found distributed throughout the BLA. Dotted lines describe the amygdalar capsule (amc) and the external capsule (ec). B) Representative fluorescence images from the mPFC a CCK-Dlx5/6-tdT. CCK^+^INs (red) are localized in all layers of the mPFC. Dotted line indicates the limit between layer 1 (L1) and layer 2. SYTO13 (green) was used to stain for cell nuclei in **A** and **B**.

Previous results have shown that hippocampal CCK^+^INs rely exclusively on presynaptic N-type calcium channels to release GABA and immunohistochemistry studies later confirmed that CCK^+^INs synapses are enriched with N-type calcium channels [12–16]. We therefore assessed if CCK^+^INs from the BLA and mPFC also rely on N-type calcium channels for GABA release. Given that CCK^+^INs synapse PNs in both the BLA and mPFC, we recorded PNs in *CCK-Dlx5/6-ReaChR.* PNs in the BLA were visually identified using DIC (**Figure 3A**). The BLA was initially visualized at 4x, and PNs were identified under 40x based on their pyramidal shape in DIC microscopy (**Figure 3A**). We confirmed PN identity based on an accommodating firing pattern recorded with current clamp, ~ 80% of recorded cells showed this firing pattern (**Figure 3B**, *left panel).* IPSC were recorded using whole-cell voltage clamp and optogenetic stimulation. This response was blocked with the GABA_A_ blocker bicuculline (BIC), demonstrating that these IPSCs rely only on GABAergic neurotransmission (**Figure 3B**, *right panel).* No blockers of excitatory transmission were used in these conditions, thus the lack of response after BIC block further supports that stimulating ReaChR results only in GABAergic neurotransmission from CCK^+^INs. To determine if N-type calcium channels contribute to GABA release from CCK^+^INs onto PNs in the BLA applied ω-ctx GVIA to the slice preparation after 20-30 min of stable IPSC recordings. Steady state of inhibition of the IPSCs inhibition by ω-ctx GVIA was reached in ~20 min (**Figure 3C**, *middle panel*). After ω-ctx GVIA block, BIC was applied to confirm that only IPSCs were being recorded (**Figure 3C**, *left and middle panels).* We found that ω-ctx GVIA ihibits ~ 50% of the IPSC in these conditions, suggesting that CCK^+^IN synapses on PNs partially rely on N-type calcium channels.

**Figure 3.**
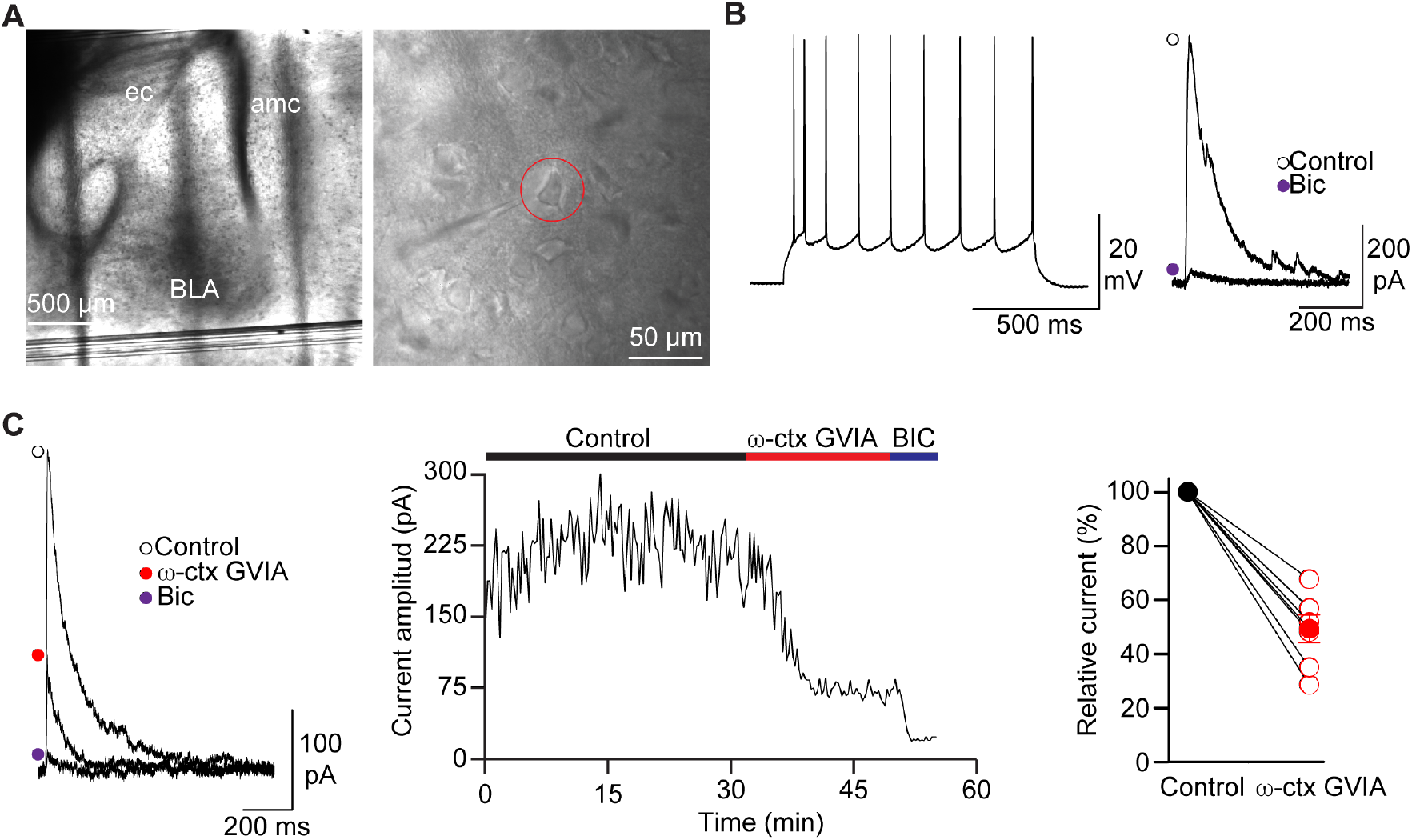
Synapses from CCK^+^INs in BLA partially rely on N-type calcium channels to release GABA. **A**) Visual identification of PNs in the BLA with DIC microscopy at 4x (left panel) and 40x (right panel). Note the pyramidal shape of the recorded cell (red circle). ec = external capsule, amc = amygdala capsule. **B**) Nonaccommodating firing pattern characteristic of PNs in the BLA (left panel). Blockage of light-evoked IPSCs with 100 μM BIC in PNs from brain slices of CCK-Dlx5/6- ReaChR mouse (right panel). **C**) Representative IPSCs with 0.5 μM ω-ctx GVIA and 100 μM BIC (left panel). Time course of ω-ctx GVIA and BIC block (middle panel). Quantification of IPSC inhibition by ω-ctx GVIA (right panel). IPSC size was determined by fitting current size to a line for 10 min of stable recordings during control and during ω-ctx GVIA exposure. Percentage of current was calculated relative to the control condition. Filled circles represent average % of current relative to control and empty circles represent % of current relative to control for each individual cell.

CCK^+^INs are also abundant in the mPFC, where they are the most common type of interneuron [49]. We next tested if CCK^+^INs in mPFC rely on N-type calcium channels to release GABA. Here, we recorded PNs of the mPFC in slices of CCK-Dlx5/6-ReaChR mice and evoked IPSCs using optogenetics. PNs were identified with DIC. We recorded PNs from layers 2/3 and 4 because they receive abundant synaptic connections from CCK^+^INs [45]. PNs in layers 2/3 and 4 were visually identified due to their pyramidal shape and the transversal orientation of their dendritic tree (towards layer 1) (**Figure 4A**). Furthermore, we identified PNs based on their characteristic non-accommodating firing pattern (**Figure 4B**, *left panel*). We recorded IPSCs in PNs, which were confirmed by full BIC block (**Figure 4B**, *right panel).* After 15 – 20 min of stable IPSC recording, ω-ctx GVIA was applied to the bath, and no significant block of the IPSC was observed during 25-30 min of recording after baseline (**Figure 4C**, *left and middle panels*). To ensure drugs were being properly perfused into the brain slice, we applied BIC, which induced full block of IPSCs (**Figure 4C**, *middle panel).* The lack of ω-ctx GVIA inhibition of IPSCs was observed in ~85% of recorded cells (**Figure 4C**, right panel), although the other 15% showed <10% block. However, on average ω-ctx GVIA failed to reduce IPSCs in CCK^+^IN synapses onto PNs of the mPFC. These results suggest that these synapses rely very little on N-type calcium channels to release GABA, which is in contrast with those results observed in CCK^+^IN synapses of the BLA.

**Figure 4.**
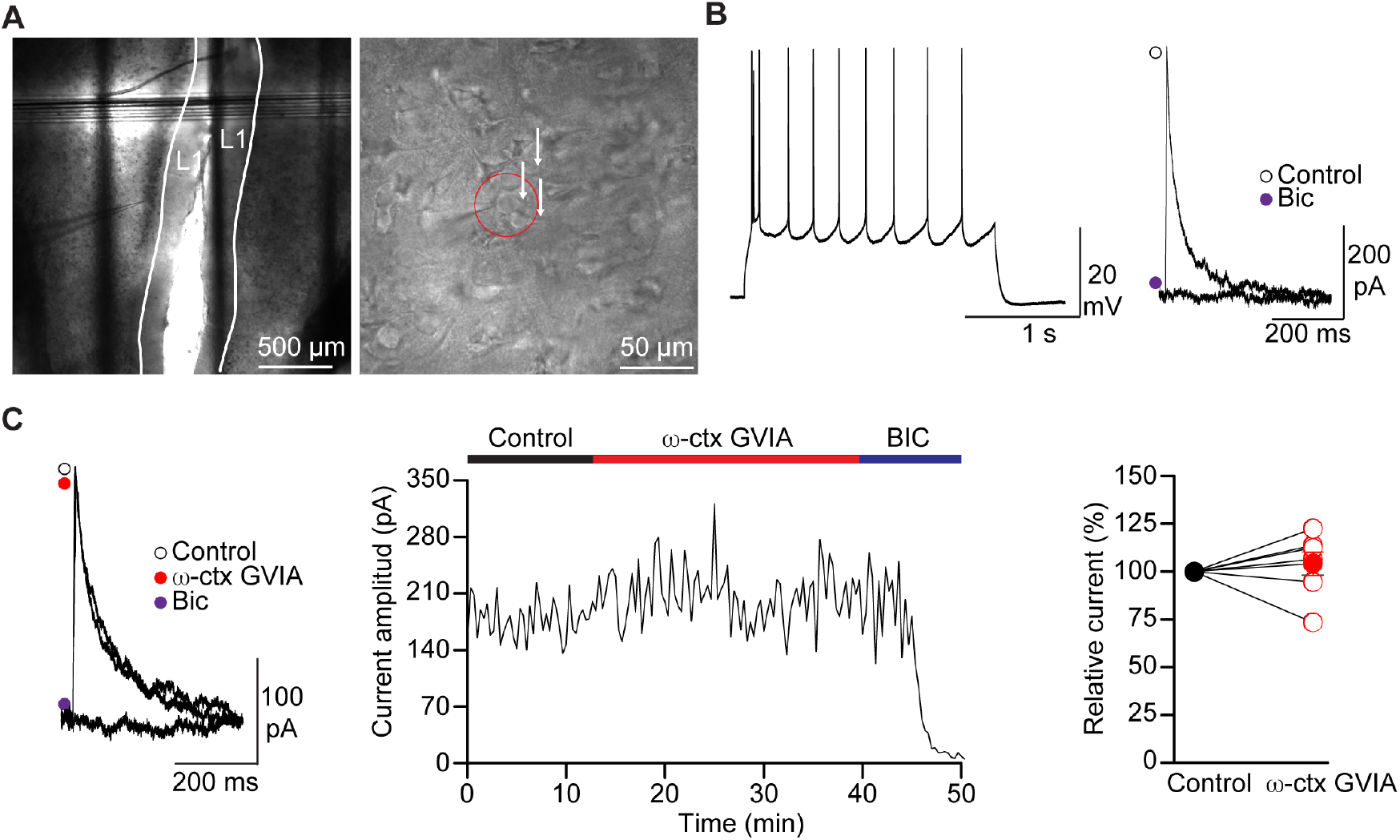
Synapses of CCK^+^INs onto PNs in the mPFC do not rely on N-type calcium channels to release GABA. **A**) Visual identification of PNs in mPFC layer 1 (L1) with DIC microscopy at 4x (left panel) and 40x (right panel). Note the pyramidal shape of the recorded cell (red circle) and the transversal orientation of dendrites arising from PNs (right panel, white arrows). **B**) Non-accommodating firing pattern characteristic of PNs in mPFC (left panel) and pharmacological characterization with 100 μM BIC of IPSCs recoded in PNs and evoked with red light in brain slices of CCK-Dlx5/6-ReaChR mouse (right panel). **C**) Representative IPSCs with 0.5 μM ω-ctx GVIA and 100 μM BIC (left panel). Time course of ω-ctx GVIA and BIC block (middle panel). Quantification of IPSC inhibition by ω-ctx GVIA (right panel). Relative current was determined as indicated above. Filled circles represent average % of current relative to control and empty circles represent % of current relative to control for each individual cell.

We next tested if GABA release from other interneurons relies on N-type calcium channels. We recorded IPSCs in PNs with electrical stimulation using a concentric electrode placed in layer I (**Figure 5A**). PNs were visually identified through their pyramidal shape and transversal dendritic orientation (**Figure 5A**, *right panel).* Similarly, we recorded a non-accommodating firing pattern. We further confirmed electrically induced IPSCs with block by BIC. In these recordings excitatory transmission was blocked with APV and NBQX, NMDA and AMPA/Kainate receptor blockers, respectively (**Figure 5B**). Full block of the IPSC by BIC in the presence of APV and NBQX suggest that GABA responses were successfully isolated in our experimental conditions. To assess if N-type calcium channels control GABA release, we applied ω-ctx GVIA to the bath after 10 – 15 min of stable IPSC recordings, we observed ~ 40% of IPSC reduction in the presence of this toxin (**Figure 5C**, *middle panel).* Inhibition by ω-ctx GVIA was observed in the vast majority of the recorded cells (**Figure 5C**, *right panel).* Our results show that inhibitory synapses onto PNs rely on N-type calcium channels. Taken together with our previous results, these synapses are likely to belong to non-CCK^+^INs.

**Figure 5.**
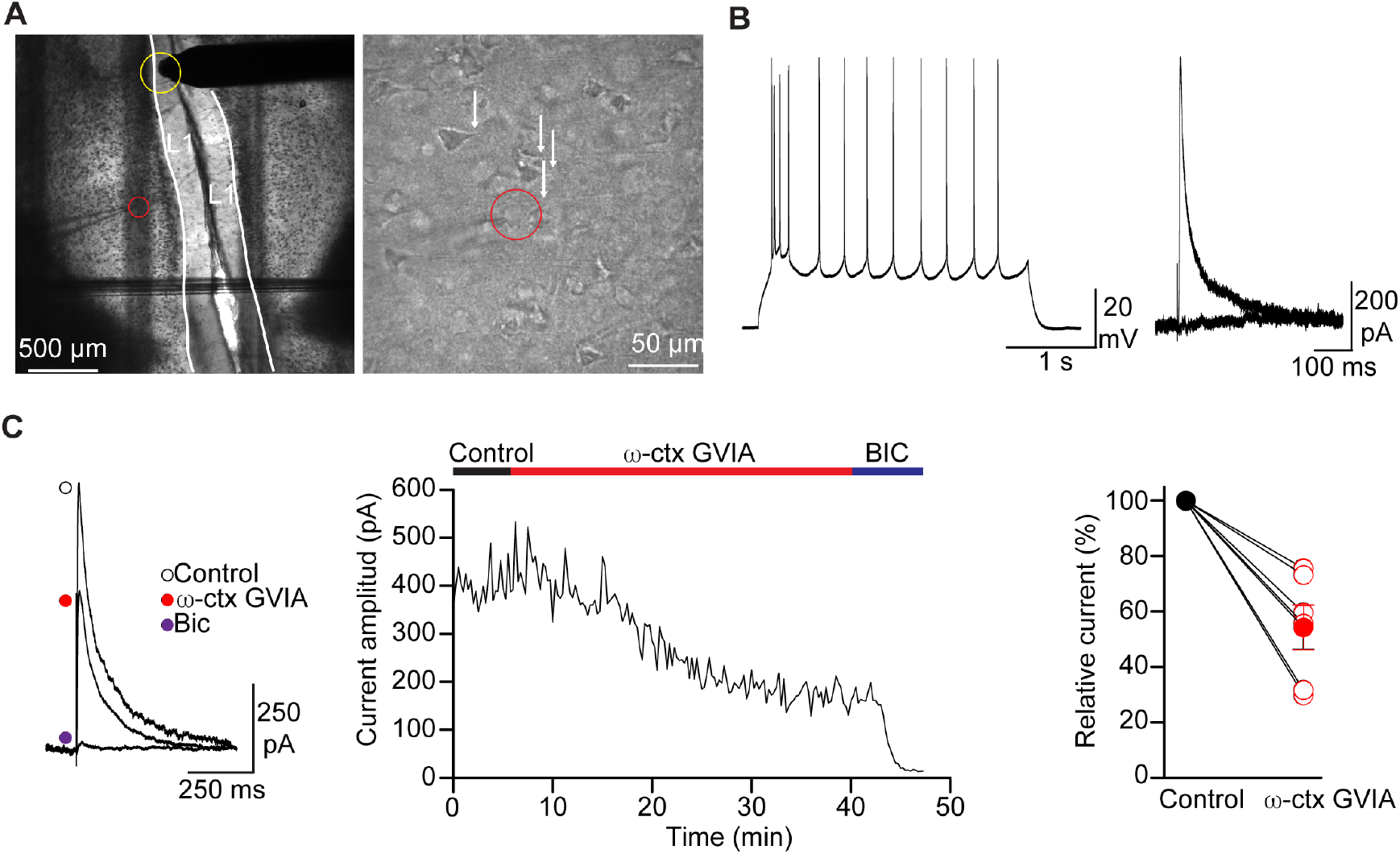
GABAergic neurotransmission in the mPFC partially relies on N-type calcium channels. **A**) Visual identification of PNs in layer 1 (L1) of the mPFC with DIC microscopy at 4x (left panel) and 40x (right panel). Note the pyramidal shape of the recorded cell (red circle) and the transversal orientation of dendrites arising from PNs (right panel, white arrows). Stimulation electrode was placed in L1 as indicated by yellow circle. **B**) Non-accommodating firing pattern characteristic of PNs from the mPFC (left panel), and pharmacological characterization with 100 μM BIC of IPSCs recoded in PNs evoked with electrical stimulation in a WT mouse (right panel). **C**) Representative IPSCs with 0.5 μM ω-ctx GVIA and 100 μM BIC (left panel). Time course of ω-ctx GVIA and BIC block (middle panel). Quantification of IPSC inhibition by ω-ctx GVIA (right panel). Percentage of current was determined as indicated previously. Filled circles represent average % of current relative to control and empty circles represent % of current relative to control for each individual cell.

## 4. Discussion

CCK^+^INs were labeled for histochemistry and electrophysiology using an intersectional labeling approach. Our results are similar to previous studies before by our group and others [27, 41, 45, 48, 49]. We found that IPSCs recorded on CCK^+^IN/PN synapses in the BLA are partially sensitive to ω-ctx GVIA. This is in contrast with previous observations in synapses of CCK^+^INs onto CA1 pyramidal cells of the HPC, where ω-ctx GVIA blocks 100% of the IPSC [12, 16]. The dominant role of N-type calcium channels on GABA release has been observed in basket CCK^+^INs and Schaffer collateral associated CCK^+^INs [13, 16]. In the BLA, CCK^+^INs are classified into CCK large and small. Large CCK^+^INs synapse onto the cell bodies of PNs that project to the mPFC and they resemble basket CCK^+^INs of the HPC [37, 51]. BLA small CCK^+^INs co-express VIP, but their function is not known [37, 51]. These observations open the possibility that different CCK^+^INs in the BLA express different proportion of presynaptic N-type calcium channels. Furthermore, it is also possible that CCK^+^INs express different amount of presynaptic calcium channels relative to CCK^+^INs in the HPC. Our results unveil a cell-specific role of N-type calcium channels in controlling transmitter release from CCK^+^INs located in brain areas linked to anxiety.

In the mPFC, we found that N-type calcium channels play a negligible role in controlling GABA release from CCK^+^INs compared to the BLA and HPC. This observation further supports a cell- and region-specific role of N-type calcium channels in the control of excitability of areas linked to anxiety. Similar to the BLA, CCK^+^INs in the cortex are heterogenous; five CCK^+^INs populations have been identified using single-cell transcriptomic analysis coupled to electrophysiology [52]. This heterogeneity could help explain the differences in the dependence on N-type calcium channels of GABA release between the BLA, mPFC and HPC. Our results align with single-cell transcriptome analysis that shows that the *Cacna1b* mRNA is expressed at low levels in only one of the five subtypes of cortical CCK^+^IN [52]. However, when using electric stimulation and blocking all glutamatergic neurotransmission, we found that the IPSCs recorded in PNs of the mPFC were partially sensitive to ω-ctx GVIA. This observation suggests that there are other types of interneurons outside of CCK^+^INs that utilize N-type calcium channels to release GABA. Accordingly, previous studies have shown that 80% of the IPSC generated by action potentials in neurogliaform interneurons of rat PFC are blocked with ω-ctx GVIA [53]. But, PV^+^INs utilize P/Q-type channels rather than N-type calcium channels [40]. This provides further support to the idea that N-type calcium channel are expressed in specific synapses of the mPFC, excluding CCK^+^IN and PV^+^IN synapses. VIP^+^INs and calbindin expressing interneurons are also present in the mPFC, however the type of presynaptic calcium channels that these interneurons use to release GABA is unknown. Further research will be needed to determine if these type of interneurons express N-type calcium channels in the mPFC.

## 5. Conclusions

In this report, we have successfully labeled CCK^+^INs using an intersectional approach. We recorded synapses of CCK^+^INs onto PNs of the BLA and mPFC. Our results show that the synaptic responses evoked with optogenetics are purely GABAergic. We found that IPSCs from CCK^+^INs onto PNs of the BLA are sensitive to ω-ctx GVIA but not fully blocked, suggesting that these synapses partially depend on N-type calcium channels to release GABA. We also showed that CCK^+^IN synapses onto PNs of the mPFC are resistant to ω-ctx GVIA, these results suggest that these synapses do not rely on N-type calcium channels to release GABA. However, we still found that other GABAergic synapses in the mPFC depend on N-type calcium channels to some degree. Given the prime role of CCK^+^INs in emotional processing, our results shed light on the possible links between N-type calcium channels and anxiety.

## Acknowledgements

We thank Melanie Bertolino and Marie Akiki for their assistant in mouse genotyping.

## Author Contributions

MB performed the histochemistry and electrophysiological experiments. BC and AB maintained, genotyped and provided input on experimental design. NC, SM, and LL helped with genotyping and maintenance of the mouse colony. NC, SM, LL, and AA wrote the manuscript. AA designed the study.

## Funding

This work was supported by the National Institute of Mental Health [grant number, R00MH099405].

## Competing Interests

Authors declare no competing interests.

## Notes

### Competing Interest Statement

The authors have declared no competing interest.

